# Heteroresistance in *Enterococcus faecalis* is prevalent for key antibiotics and mainly caused by mutations

**DOI:** 10.64898/2026.07.14.738184

**Authors:** Sheida Heidarian, Sarah Lohsen, Andrei Guliaev, Hervé Nicoloff, Sarah Satola, Dan I. Andersson, Karin Hjort

## Abstract

Heteroresistance (HR), the coexistence of a rare resistant subpopulation within a predominantly susceptible bacterial population, is a clinically relevant problem. The resistant subpopulation often escapes detection by standard susceptibility testing, which can lead to treatment failure. To investigate HR in *Enterococcus faecalis*, we performed population analysis profiling (PAP) on 40 clinical isolates against five clinically important antibiotics and whole-genome sequenced (WGS) the resistant subpopulations. HR was identified for daptomycin (20.0%), gentamicin (13.2%), and tigecycline (35.9%), but not for linezolid, and vancomycin. Genomic analysis revealed that daptomycin resistance was primarily caused by mutations affecting cell envelope integrity and stress-response pathways. Gentamicin resistance was linked to alterations in efflux regulation, ribosomal proteins synthesis, and transcriptional control. Tigecycline resistance involved deletions in the *tet*(M) leader peptide, resulting in a 25-fold increased *tet*(M) expression, as well as transposition of the transposon Tn916 carrying *tet*(M) to multiple chromosomal sites, causing an increased gene dosage of *tet*(M). These findings highlight the role of chromosomal mutations and mobile genetic elements in causing HR in *E. faecalis*

**Author Summary:** Antibiotic resistance is one of the most critical challenges in modern medicine, undermining the efficacy of antimicrobial therapies and compromising patient safety. Bacterial strains that are heteroresistant (HR), show a minor subpopulation of resistant bacteria within a main susceptible bacterial population. HR strains are mostly classified as susceptible strains since the subpopulation are usually too small to be detected with standard susceptibility testing. Treatment with antibiotics will lead to survival and increase of the resistant subpopulation that can lead to treatment failure. In this study, we observed a high frequency of HR among clinical *Enterococcus faecalis* strains against gentamicin, daptomycin and tigecycline, three commonly used antibiotics for treatment of *E. faecalis* infections. Chromosomal mutations and changes in gene expression were the genetic mechanisms generating the resistant subpopulations.

## Introduction

Antibiotic resistance represents one of the most critical challenges in modern medicine, undermining the efficacy of antimicrobial therapies and compromising patient safety worldwide. Among the mechanisms of resistance, heteroresistance (HR) has gained increasing attention as a clinically significant phenotype (1). HR is defined as the presence of a minor resistant subpopulation within a susceptible main population. Under antibiotic selective pressure, the resistant subpopulations can expand and thereby cause treatment failure (1–4). In HR isolates, resistant subpopulations occur at low frequencies (typically ≈ 10^-7^ to 10^-4^) and because of their transient phenotype and low frequency they are often misclassified as susceptible using conventional diagnostic tools (1,5,6).

HR has been documented for multiple bacterial species and antibiotic classes (6–9). The gold standard method for HR detection is population analysis profiling (PAP), and HR is typically defined as the presence of subpopulations (≥10⁻⁷ frequency) with a minimum inhibitory concentration (MIC) at least 8-fold higher than the MIC of the main population (4,7). However, differences in HR definitions and the labor-intense nature of the PAP analysis limit its use in the clinic, leading to uncertainties regarding HR prevalence in clinical settings (6). Several studies have demonstrated the contribution of these resistant subpopulations to antibiotic therapy failure, prolonged infections, and the emergence of fully resistant strains, both in animal models and clinical settings (10–15).

At the molecular level, HR arises through diverse genetic mechanisms, including genetic alterations via point mutations, insertions, deletions, and increased gene-dosage (1,8,9,16,17). The latter can be the result of tandem gene amplifications, increased plasmid copy number, or mobilization of resistance genes onto high-copy plasmids or into highly expressed genes. In Gram-negative bacteria, tandem gene amplification and plasmid copy number increases are recognized as the dominant HR mechanisms (9,17–20). In contrast, for Gram-positive bacteria, HR more frequently arises from genetic alterations such as mutations or deletions (8), although amplification-based mechanisms have also been reported for vancomycin (21,22).

The majority of HR studies in Gram-positive bacteria have focused on *S. aureus*, particularly the heteroresistant vancomycin-intermediate *S. aureus* (hVISA) phenotype (23,24). More recently, HR has also been described for additional antibiotic classes such as β- lactams, aminoglycosides, lipopeptides, and glycopeptides (8,25,26). A few studies have also investigated HR in other Gram-positive pathogens, including *Streptococcus* and *Enterococcus*. Among enterococci, *Enterococcus faecium* and *Enterococcus faecalis* are major opportunistic pathogens and ranked second only to *Staphylococcus* as the cause of hospital-acquired infections (27–29). These species display remarkable adaptability to diverse environments and possess both intrinsic and acquired antimicrobial resistance mechanisms, making them an increasing clinical concern. Indeed, the World Health Organization has classified vancomycin-resistant *E. faecium* as a high-priority pathogen requiring urgent development of new therapies (30). Most reports on enterococcal HR focus on *E. faecium* (31–33) and particularly glycopeptide resistance (21,33) whereas studies on *E. faecalis* HR remain limited. However, recent reports, have documented HR in *E. faecalis* against omadacycline (34) and eravacycline (35). Understanding the prevalence and underlying mechanisms of HR in *E. faecalis* is crucial for several reasons. First, undetected HR can contribute to treatment failure. Second, the molecular basis of HR in enterococci often involves plasmid-borne resistance genes, which can undergo amplification, overexpression, or increases in plasmid copy number (21,22,34,36). This mechanism partly differs from *S. aureus*, where chromosomal point mutations are more frequently responsible for HR (8). Third, both HR (31) and non-HR (37) enterococcal isolates exhibit a strong capacity to acquire and disseminate plasmid-mediated resistance determinants, underscoring their role in antimicrobial resistance (AMR) spread. Fourth, most HR studies in enterococci have focused on *E. faecium* and on specific antibiotics, leaving the prevalence and mechanisms of HR toward other agents poorly understood. Finally, the full clinical relevance of HR remains uncertain, with only a few cases linking HR to adverse patient outcomes and these exclusively in *E. faecium* (38,39). These observations highlight the importance of investigating HR in *E. faecalis* to elucidate its role in treatment failure, prolonged infection, and the wider AMR crisis.

In this work, we determined the prevalence of HR and identified mutations that can cause HR among 40 clinical *E. faecalis* isolates tested against different clinically relevant antibiotic classes by using a consistent method and HR definition. We demonstrate that: (i) the prevalence of HR varied widely among different antibiotic classes; (ii) the frequency of clinical breakpoint (CBP) crossing HR (subpopulations growing above the CBP) was notable for some antibiotics; (iii) chromosomal point mutations, deletions in gene regulatory region resulting in enhanced gene expression, and increased gene dosage due to transposon-mediated insertions were the predominant mechanisms generating the resistant subpopulations and the HR phenotype.

## Material and Methods

### Bacterial isolates, growth conditions, and chemicals

A total of 40 clinical *E. faecalis* strains were collected from four European countries (Sweden, Spain, Denmark, and Norway; 10 isolates per country). Five clinically relevant antibiotics were tested: daptomycin (DAP), gentamicin (GEN), linezolid (LNZ), tigecycline (TGC), and vancomycin (VAN). All antibiotics were purchased from Sigma-Aldrich, except for tigecycline (Tygacil, Wyeth), which was purchased from Apoteket. Unless otherwise specified, isolates were cultured at 37°C in Mueller Hinton broth II (MHB II) with shaking at 220 rpm or on Mueller Hinton II agar (MHA II) (BD, Germany). Antibiotic stock solutions were prepared fresh prior to use, and antibiotic-containing agar plates were prepared immediately before each experiment.

### Antimicrobial susceptibility tests

Antimicrobial susceptibility to DAP, GEN, LNZ, TGC, and VAN was determined using Etest gradient strips (bioMérieux, Marcy l’Étoile, France) following the manufacturer’s instructions. The MIC and the clinical break point were interpreted according to the European Committee on Antimicrobial Susceptibility Testing (EUCAST) 2025 guidelines. Since clinical breakpoints for DAP and GEN are not available, isolates were characterized using the epidemiological cutoff value (ECOFF) from EUCAST.

### Population Analysis Profiling (PAP) tests

PAP tests were performed using a modified microdilution plating protocol (19,40). Strains were streaked from frozen glycerol stocks onto MHA II and incubated overnight at 37°C. Single colonies were inoculated into 48-well plates containing 300 µl MHB II and cultured for ∼20 h at 37°C with shaking (500 rpm). Overnight cultures were serially diluted (10-fold, seven steps) in MHB II, and 7.5 µl of each dilution was spotted onto MHA II without antibiotic for colony-forming unit (CFU) enumeration. For DAP, dilutions were also spotted onto MHA (II) supplemented with 2–32 mg/L (two-fold increments). In contrast, for GEN, LNZ, TGC, and VAN, 100 µl of overnight culture was spread over the full surface of MHA II plates containing the clinical breakpoint concentration as well as 4× and 8× the MIC of each isolate using glass beads. Antibiotic concentrations tested were GEN (32–128 mg/L), LNZ (6–24 mg/L), TGC (0.064–1.52 µg/ml), and VAN (4–24 mg/L). Plates were incubated at 37°C, and final CFU counts were recorded after 40–48 h. The HR phenotype was defined as subpopulation growth at concentrations ≥8-fold above the MIC of the main population with a frequency of 10⁻⁷ or above. Isolates meeting this criterion were selected for further mechanistic analysis.

### Selection of Resistant Mutants

Resistant mutants were selected for parental isolates exhibiting the HR phenotype at 8× above the MIC (Supplementary Table 1). Overnight cultures were prepared in MHB II, and approximately 10⁸ cells were plated on MHA II containing 8× MIC of the corresponding antibiotic. Colonies appearing after 48 h were picked, re-streaked on MHA II with the same antibiotic concentration, and inoculated into 21 ml MHB II supplemented with the same antibiotic concentration. After overnight incubation, 1 ml of each culture was frozen in 10% DMSO at -80°C for long-term storage, while the remaining 20 ml was pelleted and stored at - 20°C for DNA extraction and WGS. The resistance level of each selected mutant was determined by Etest, following the same methodology and interpretive criteria described in the antimicrobial susceptibility testing section.

### Whole genome sequencing

WGS was performed on both the parental HR isolates and selected resistant clones. Genomic DNA was extracted using the MasterPure Gram Positive DNA Extraction Kit (Lucigen). For parental isolates, DNA was obtained from 1 ml overnight cultures, while 20 ml cultures were used for resistant mutants to achieve adequate amount of cell pellet, as they were slow growing. To improve cell lysis for the standard protocol, two modifications were introduced: (i) an addition of 2 μl lysostaphin (50 mg/ml) (Sigma-Aldrich) per ml of pelleted cell culture and an extended digestion time of 45 min, and (ii) an addition of 4 μl proteinase K per each DNA extraction (Sigma-Aldrich) to enhance digestion efficiency. Parental isolates were sequenced using both Nanopore (MinION Mk1C, rapid barcoding kit 96, multiplexing up to 40 genomes per R9 cell) and DNBseq (800 bp paired-end libraries, BGI, Warsaw). Nanopore long reads were initially filtered for a minimum length of 3,000 nucleotides using filtlong v0.2.1 (41), and adapters were trimmed with porechop v0.2.4 (42). Resistant clones were sequenced with DNBseq only. Short-read quality was controlled using fastqc v0.11.9 (43) and 10 nucleotides from the 5’-end of each read was trimmed using fastp v0.20.1 (44). Genome assemblies were generated using two different hybrid approaches: a combination of flye v2.9.1 (45), medaka v1.6.0 and polypolish v0.5.0 (46) was used for assembling the genomes when long read coverage was higher than 20X, otherwise Unicycler v0.4.8 (47) was used. Assembly quality was assessed using QUAST v5.0.2 (48) and BUSCO v5.2.2 (49). Genome annotation was performed with Prokka v1.14.6 (50). All analyses were automated and made reproducible using Snakemake v7.32.4 (51). DNBseq reads from resistant clones were mapped to their parental genomes using CLC Genomic Workbench (Qiagen), and variants (SNPs, indels, structural variants, and amplifications) were identified through coverage and variant detection tools. Resistance genes were detected with the CARD database (strict cutoff: >90% identity and >90% coverage) (52)Repeat regions within 200 kbp of resistance genes were manually inspected in CLC. Sequencing data are deposited in NCBI under the BioProject PRJNA1452924.

### RNA extraction and quantitative Real-Time PCR (qRT-PCR)

Total RNA was extracted from TGC resistant mutants and their parental HR isolates. Three independent overnight cultures per isolate were grown in brain heart infusion (BHI) broth, diluted 1:50 into fresh medium, and incubated at 37°C with shaking until reaching mid-exponential phase (OD₆₀₀ ∼ 0.3). RNAs were stabilized with RNAprotect Bacteria Reagent (Qiagen) and bacterial cells were pelleted by centrifugation. RNA was purified using the RNeasy Mini Kit (Qiagen) with modified lysis conditions: pellets were treated with lysozyme (15 mg/mL) and proteinase K (100 µg/mL) in TE buffer for 10 min, followed by lysis in RLT buffer supplemented with β-mercaptoethanol. RNA was column-purified according to the manufacturer’s protocol and eluted in RNase-free water. Residual genomic DNA was removed using the TURBO DNA-free Kit (Ambion). cDNA was synthesized using the High-Capacity cDNA Reverse Transcription Kit (Applied Biosystems), including negative controls lacking RNA or reverse transcriptase. Gene expression was quantified by SYBR Green-based qRT-PCR in 20 µL reactions containing 2× SYBR Supermix, primers (1 µM each), nuclease-free water, and 1 µL diluted cDNA. Cycling conditions were 50°C for 2 min, 95°C for 10 min, followed by 40 cycles of 95°C for 15 s, 57°C for 30 s, and 72°C for 30 s. Primer sets targeting the *tet*(M) gene (gene of interest), *gdh* (housekeeping gene) (53,54), was used (Supplementary Table 2). Amplification efficiency was verified using standard curves from serial dilutions, and no-template controls were included. Gene expression of *tet(*M) normalized to *gdp* and calculated by the ΔΔCt method and relative expression levels were calculated as fold-change of resistant mutants compared to their respective parental isolates. Statistical significance was assessed by Student’s *t*-test.

### Cross-resistance and collateral sensitivity

To investigate cross-resistance (CR), the MIC levels of the resistant mutants selected from the HR strains were determined towards seven antibiotics. Bacterial cells were recovered by scraping approximately 30 µL of culture from frozen stocks and suspending them in saline to achieve a turbidity equivalent to a 0.5 McFarland standard. The suspension was evenly spread onto MHA II plates using sterile cotton swabs. MICs were determined for each individual isolate using Etest strips toward antibiotics not used in the initial selection of the respective resistant mutant, following the same culturing conditions and interpretation criteria as described above for antimicrobial susceptibility testing. To calculate CR, the following formula was used: log2 [MIC (resistant mutant)/MIC (parental isolate)].

### Statistical analysis

The correlation between the MIC of isolates and the HR phenotype was examined using the Mann-Whitney U test. To evaluate the significancy of relative gene expression of resistant mutant to their respective parental isolate the student t-test was used. All statistical analyses were performed using GraphPad Prism (10.6.1). Two-sided tests were employed, and a P-value of 0.05 was considered as statistically significant.

## Results

### Susceptibility profiles of clinical *E. faecalis* isolates

MICs of DAP, GEN, LNZ, TGC, and VAN were determined for 40 *E. faecalis* clinical isolates (Table 1). All isolates that were classified as susceptible according to EUCAST clinical breakpoints or to the epidemiological cutoff value (ECOFF) according to EUCAST were included in the HR analysis. MIC distributions varied across antibiotics (Supplementary Fig. 1) and GEN exhibited the widest MIC range (3–32 mg/L), whereas other antibiotics showed narrower distributions. For all antibiotics except GEN, the observed MIC distributions closely aligned with EUCAST international reference datasets (20,000–30,000 isolates per antibiotic; mic.eucast.org). For GEN, the isolates demonstrated lower MICs with 75% (30/40) of the isolates having MICs between 3-6 mg/L, compared to the EUCAST reference distribution, which peaks between 8-32 mg/L. All 40 isolates had MICs below the clinical breakpoint or the ECOFF value for at least one of the antibiotics tested.

**Table 1.** Minimal inhibitory concentrations (MICs) of 40 clinical *Enterococcus faecalis* isolates against seven clinically relevant antibiotics, measured by single E-test for each isolate–antibiotic combination.

| Isolate ID | Country of origin | AMP | AMX | DAP | GEN | LNZ | TGC | VAN |
| --- | --- | --- | --- | --- | --- | --- | --- | --- |
| DA 70200 | Spain | 0.75 | 0.5 | 1.5 | 3 | 2 | 0.19 | 2 |
| DA 70202 | Spain | ND | ND | 1 | 4 | 2 | 0.064 | 1 |
| DA 70204 | Spain | ND | ND | 1 | 32 | 1.5 | 0.016 | 1.5 |
| DA 70206 | Spain | 1 | 0.75 | 0.75 | 4 | 2 | 0.032 | 1 |
| DA 70208 | Spain | ND | ND | 1 | >32 | 1.5 | < 0.016 | 1 |
| DA 70212 | Spain | ND | ND | 3 | 8 | 1 | 0.032 | 1.5 |
| DA 70214 | Spain | ND | ND | 2 | 6 | 1.5 | 0.064 | 2 |
| DA 70218 | Spain | 0.75 | 0.5 | 2 | 4 | 1.5 | 0.032 | 1 |
| DA 70220 | Spain | 0.75 | 0.5 | 1.5 | >32 | 1.5 | 0.023 | 1 |
| DA 70222 | Spain | ND | ND | 1.5 | 12 | 2 | 0.047 | 1 |
| DA 70458 | Sweden | ND | ND | 2 | 6 | 2 | 0.047 | 1.5 |
| DA 70460 | Sweden | ND | ND | 2 | 4 | 2 | 0.023 | 3 |
| DA 70464 | Sweden | ND | ND | 1.5 | 6 | 3 | 0.032 | 1 |
| DA 70466 | Sweden | ND | ND | 2 | 8 | 3 | 0.032 | 2 |
| DA 70468 | Sweden | 0.5 | 0.5 | 1 | 3 | 2 | 0.032 | 2 |
| DA 70470 | Sweden | ND | ND | 0.75 | 32 | 1 | 0.023 | 1 |
| DA 70472 | Sweden | ND | ND | 1 | 4 | 1.5 | 0.032 | 1 |
| DA 70474 | Sweden | ND | ND | 2 | 16 | 1.5 | 0.047 | 1 |
| DA 70476 | Sweden | ND | ND | 1.5 | 4 | 1 | 0.032 | 1 |
| DA 70480 | Sweden | ND | ND | 1.5 | 6 | 1 | 0.032 | 1 |
| DA 70594 | Norway | ND | ND | 1.5 | 4 | 2 | 0.064 | 2 |
| DA 70596 | Norway | 0.75 | 0.5 | 2 | 6 | 1.5 | 0.032 | 2 |
| DA 70600 | Norway | ND | ND | 1.5 | 2 | 1.5 | < 0.016 | 1 |
| DA 70602 | Norway | ND | ND | 3 | 6 | 1.5 | 2 | 1.5 |
| DA 70604 | Norway | ND | ND | 1 | 4 | 2 | 0.032 | 0.75 |
| DA 70606 | Norway | ND | ND | 0.38 | 3 | 1 | 0.023 | 1 |
| DA 70608 | Norway | ND | ND | 1 | 4 | 1 | 0.032 | 1 |
| DA 70610 | Norway | ND | ND | 1 | 4 | 1.5 | 0.032 | 3 |
| DA 70612 | Norway | 1.5 | 1.5 | 2 | 6 | 1 | 0.047 | 1.5 |
| DA 70614 | Norway | ND | 0.5 | 1 | 4 | 1 | 0.064 | 1 |
| DA 70814 | Denmark | ND | ND | 2 | 8 | 1.5 | 0.125 | 1 |
| DA 70816 | Denmark | ND | ND | 0.75 | 6 | 1 | 0.064 | 1.5 |
| DA 70820 | Denmark | ND | ND | 1 | 6 | 0.75 | 0.023 | 1.5 |
| DA 70822 | Denmark | ND | ND | 1 | 4 | 1.5 | 0.032 | 2 |
| DA 70824 | Denmark | ND | ND | 0.75 | 6 | 1.5 | 0.047 | 1 |
| DA 70828 | Denmark | ND | ND | 0.75 | 6 | 1 | 0.094 | 1 |
| DA 70830 | Denmark | ND | ND | 0.75 | 4 | 1 | 0.032 | 1 |
| DA 70832 | Denmark | 0.38 | 0.5 | 1 | 3 | 0.75 | 0.032 | 1 |
| DA 70834 | Denmark | ND | ND | 2 | 4 | 1.5 | 0.032 | 1 |
| DA 70836 | Denmark | 0.75 | 0.5 | 2 | 8 | 1.5 | 0.047 | 1 |

### Prevalence and distribution of HR in clinical *E. faecalis* isolates

To assess the prevalence of HR and identify resistant subpopulations, PAP tests were performed on the 40 *E. faecalis* clinical isolates against five antibiotics. Three isolates (two for GEN and one for TGC) were resistant to the antibiotic and not investigated. HR was defined as the presence of a resistant subpopulation at a frequency of ≥10⁻⁷ at ≥8-fold the MIC of the main population in at least one of three biological replicates. No HR phenotype was detected for LNZ or VAN. In contrast, HR was observed for DAP, GEN, and TGC at frequencies of 20.0% (8/40 isolates), 13.2% (5/38 isolates), and 35.9% (14/39 isolates), respectively (Table 2, Fig. 1). Among these, 75% (6/8) of DAP-HR and 20% (1/5) of GEN-HR isolates had subpopulations growing at antibiotic concentrations exceeding the ECOFF, and 21.4% (3/14) of TGC-HR isolates had subpopulations growing at antibiotic concentrations above the clinical breakpoint. When calculated against all isolates tested, breakpoint-crossing HR (BCHR) frequencies were 15% (6/40 isolates), 2.6% (1/38 isolates), and 7.7% (3/39 isolates) for DAP, GEN, and TGC, respectively (Table 2). Analysis of HR distribution revealed that 42.5% (17/40) of the isolates were non-HR to all five antibiotics, while 47.5% (19/40) and 10% (4/40) exhibited HR to one or two antibiotics, respectively (Fig. 1). Among 50 antibiotic–isolate combinations (five antibiotics, 10 isolates per location) tested for the HR phenotype, 20% (10/50) of HR cases toward DAP, GEN, and TGC were from Spain, 10% (5/50) from Norway, 10% from Sweden, and 14% (7/50) from Denmark. For individual antibiotics, 37.5% (3/8, DAP), 40% (2/5, GEN), and 35.7% (5/14, TGC) originated from Spain (Fig. 1). These differences are not statistically significant but might indicate country differences in prevalence. No significant MIC differences were observed between HR and non-HR isolates; for DAP and GEN, non-HR isolates had slightly higher median MICs, while TGC values were identical (Supplementary Fig. 2). MIC distributions for DAP and GEN were comparable, with each HR isolate having a non-HR counterpart with the same MIC (Supplementary Fig. 2).

**Fig. 1.**
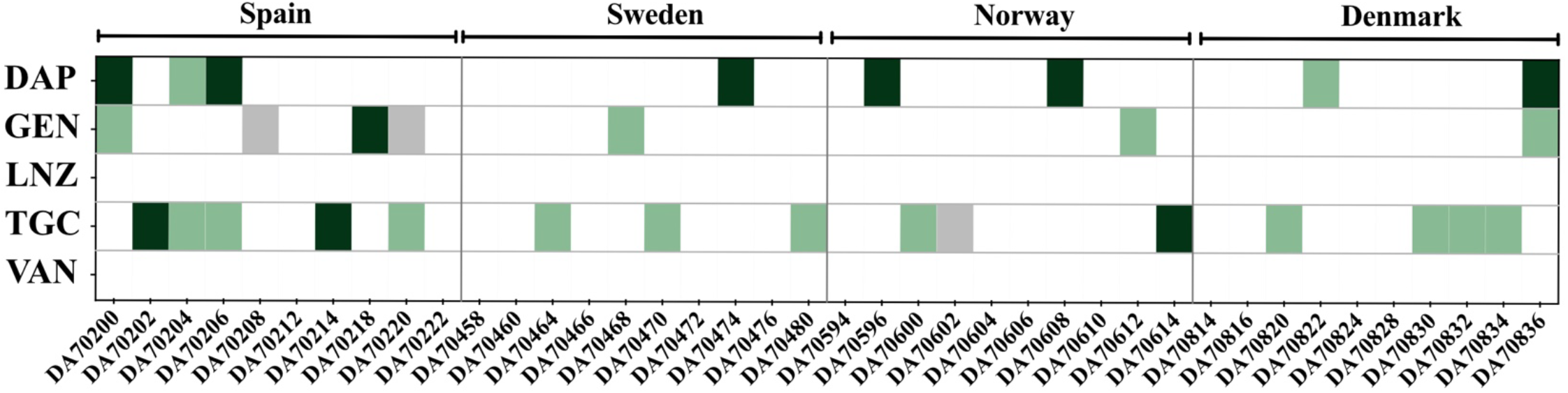
Heteroresistance (HR) profiles of 40 clinical *Enterococcus faecalis* isolates across five antibiotics. Green indicates isolates exhibiting HR as determined by population analysis profiling (PAP), while white represent isolates with non-HR phenotypes. Isolates resistant to a given antibiotic (highlighted in grey) were excluded from PAP analysis. Dark green denotes HR isolates with subpopulations exceeding the EUCAST clinical breakpoint or epidemiological cutoff (ECOFF) for the corresponding antibiotic. Isolates are grouped by country of origin, as indicated by horizontal lines above the matrix.

**Table 2.** Prevalence (% fraction of parental isolates) of heteroresistance among susceptible clinical *E. faecalis* isolates for five different antibiotics. EUCAST clinical breakpoints (CBPs) were applied for all antibiotics except daptomycin (DAP) and gentamicin (GEN), for which epidemiological cut-off values (ECOFFs) were used. CBPs: AMP (Ampicillin) >4 mg/L; AMX (Amoxicillin) >4 mg/L; LNZ (linezolid) >4 mg/L; TGC (Tigecycline) >0.5 mg/L; VAN (vancomycin) >4 mg/L; ECOFFs: DAP (daptomycin) >4 mg/L; GEN (gentamicin) >64 mg/L

| Antibiotic | Number of susceptible parental isolates analyzed | Number (percentage) of parental isolates showing HR by PAP test | Number (percentage) of parental isolates showing HR with subpopulations above EUCAST clinical breakpoint |
| --- | --- | --- | --- |
| DAP | 40 | 8 (20%) | 6 (15%) |
| GEN | 38 | 5 (13.2%) | 1 (2.6%) |
| LNZ | 40 | 0 (0%) | 0 (0%) |
| TGC | 39 | 14 (35.9%) | 3 (7.7%) |
| VAN | 40 | 0 (0%) | 0 (0%) |

### Resistance profiles of antibiotic-resistant mutants derived from HR subpopulations

Resistant mutants capable of growing at concentrations ≥8-fold higher than the MIC of the parental isolate were selected using four different parental isolates for GEN and TGC and five isolates for DAP. A total of 23 resistant mutants for DAP, 19 for GEN, and 17 for TGC were selected and analyzed. Resistance levels of the mutants differed relative to their respective parental isolates. For GEN and TGC, most mutants exhibited at least a 4-fold increase in MIC (14/19 (73.7%) and 14/17 (82.4), respectively), whereas for DAP, only 8 out of 23 (34.8%) mutants displayed a similar ≥4-fold MIC increase (Supplementary Fig. 3).

### Three distinct genetic mechanisms underlying DAP, GEN, and TGC resistance in HR strains

To examine the mechanisms underlying resistance in HR strains, we performed WGS of 59 resistant mutants selected at 8x above MIC of the parental isolate with DAP (23 isolates), GEN (19 isolates), and TGC (17 isolates), along with their 11 corresponding parental isolates. A majority of the resistant mutants carried a single genetic alteration, although GEN mutants more frequently acquired two mutations (32%, 5 isolates) (Supplementary Fig. 4A–C).

In all of the 23 analyzed DAP-resistant mutants, diverse types of mutations, including amino acid substitutions, frameshift mutations, deletions, and inversions were identified and they were consistently associated with at least a two-fold increase in MIC values compared to their respective parental isolates (Figs. 2, 3 and, Supplementary Fig. 4A). Exceptions were observed for four mutants, which carried mutations but had an identical MIC to that of their parental isolates. A majority, 78% (18/23), of the DAP resistant mutants harbored mutations in *liaX*, *liaF*, or *liaS*, genes of the *liaFSR* regulatory system involved in cell envelope stress response in enterococci (55,56). Four of these isolates shared an identical three-nucleotide insertion in *liaF*, resulting in a leucine 171 duplication. Three additional mutants carried alterations in a DUF4097 family β-strand repeat-containing protein, a gene not previously linked to DAP resistance (Supplementary Fig. 4A).

**Fig 2.**
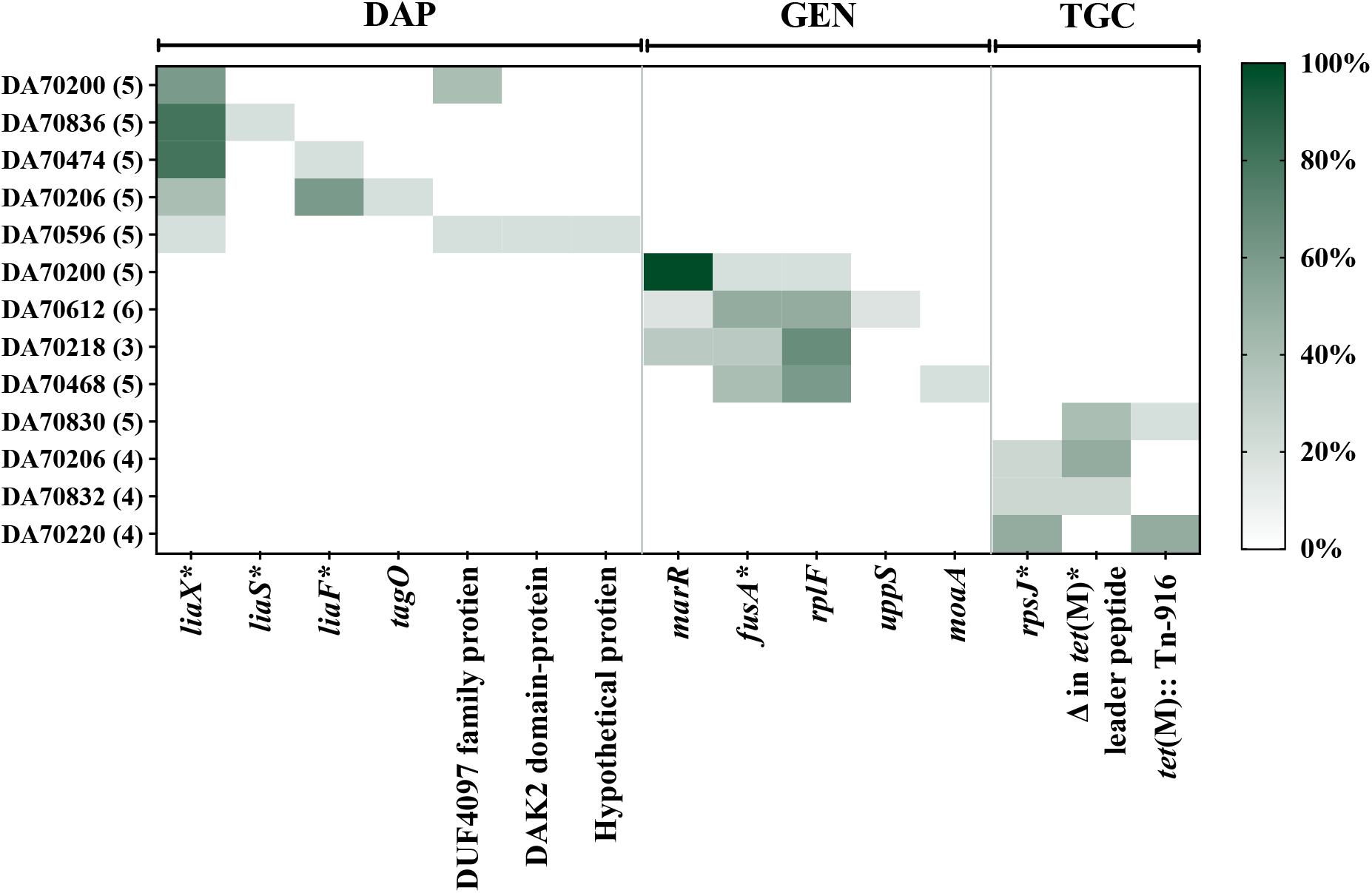
Frequency of mutations in heteroresistant isolates. The x-axis denotes gene names, and the y-axis lists the parental isolates, with the total number of mutants derived from each parental strain indicated in parentheses. Genes known to confer resistance to the specific antibiotic when mutated in *Enterococci* are marked by a asterisk (*). All mutations present in each isolate are presented in supplementary Fig. 4. The light-to-dark green scale represents the frequency of resistant mutants carrying a mutation in the same gene.

For GEN, all resistant mutants carried mutations in at least one out of the three genes: *marR*, *fusA*, or *rplF* (Figs. 2, 3 and, Supplementary Fig. 4B). The most frequently mutated locus was *rplF* (47.4%, 9/19 mutants), followed by *fusA* and *marR* found in 36.8% (7/19) of the mutants each (Supplementary Fig. 4B). Mutations in all three genes are known to contribute to GEN resistance in different species of bacteria (57), but none of the above genes have previously been reported to be involved in resistance in Enterococcus (61,62). Two different mutations were observed in 32% (6/19) of the mutants (Supplementary Fig. 4B). Of these six isolates, three carried mutations in both *marR* and *fusA*, one exhibited mutation in *marR* and *rplF*, and the remaining two possessed a mutation in either *fusA* or *rplF* in combination with a mutation in *moaA* or *uppS.* As *moaA* and *uppS* are not linked to aminoglycoside resistance, except for the reported role of *uppS* in low-level bacteriocin resistance in *E. faecalis* (58), their contribution might be negligible given the presence of mutations in established resistance genes (*fusA* and *rplF*).

Among 17 TGC-resistant *E. faecalis* mutants, four harbored mutations in *rpsJ* encoding the ribosomal S10 protein, a known determinant of TGC resistance (59,60) (Figs. 2 and 3, Supplementary Fig. 4C). Three isolates had an identical A54E substitution, and one had a 12-bp deletion. In addition, eight mutants were altered in *tet*(M), either through deletions in the leader peptide region (n = 5) or via multiple chromosomal insertions of the transposon Tn916 carrying *tet*(M), which integrated at 6 to 12 different chromosomal sites (n = 3; Fig. 4B). In five mutants with deletions within the leader peptide 40 nucleotides upstream of *tet*(M), RT-qPCR was performed to assess the impact of the mutations on expression levels of *tet*(M). Gene expression of *tet*(M) normalized to *gdp* revealed a significant upregulation of *tet*(M) (∼25-fold; Student’s t-test, p <0.001) (Fig. 4A) as compared to the parental isolate. Five mutants showed no detectable genetic changes by WGS, and the resistance mechanism in these remains undetermined.

**Fig 3.**
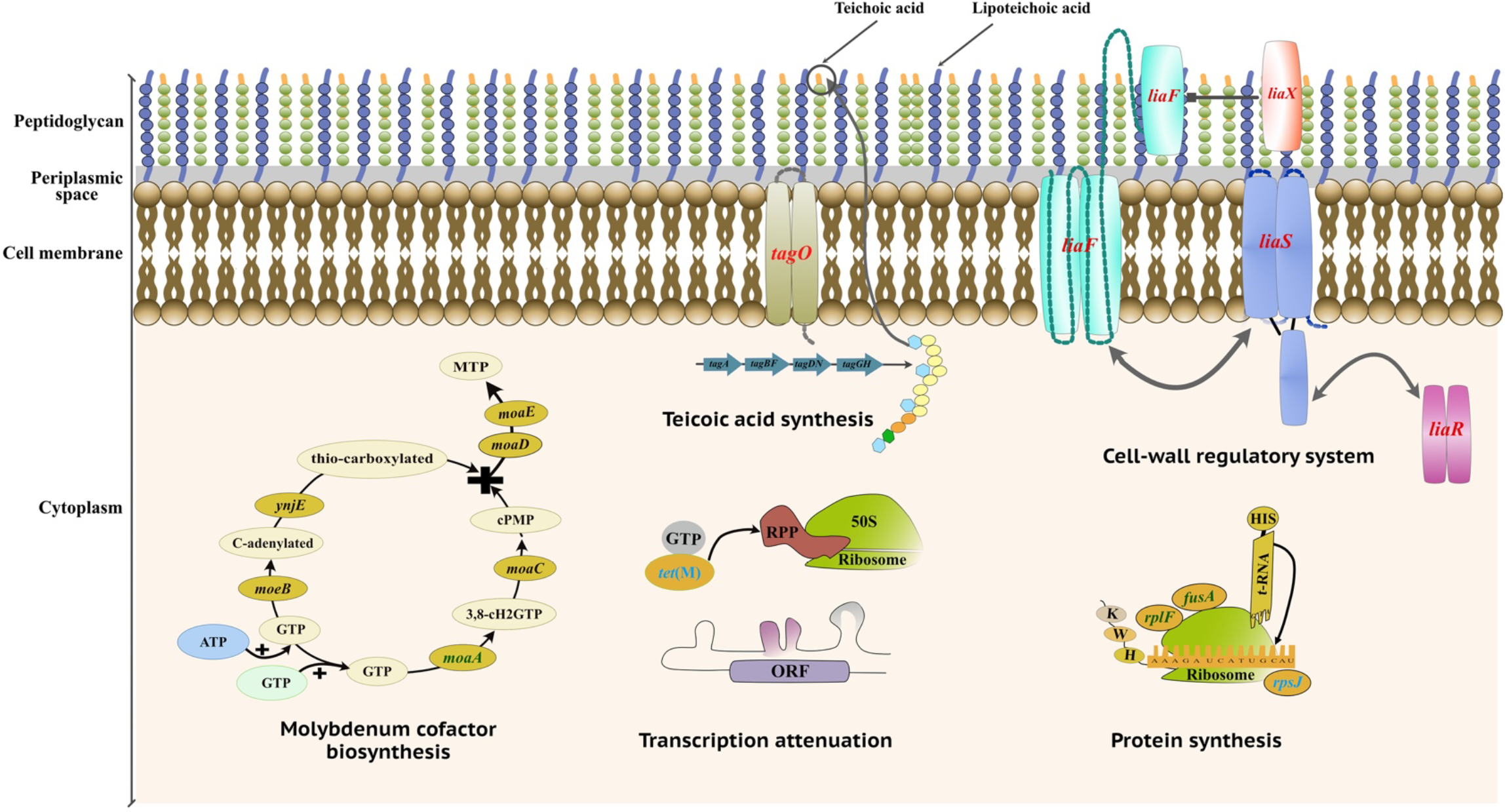
Schematic representation of mutated pathways in resistant mutants. The mutated genes, highlighted in red for DAP, green for GEN, and blue for TGC, identified in DAP, GEN, and TGC resistant mutants were involved in a diverse range of pathways, including peptidoglycan synthesis (*tagO*); protein synthesis (*rpsJ*, *rplF*, and *fusA*); molybdenum cofactor biosynthesis (*moaA*); cell wall regulatory system (*liaF*, *liaS*, *liaR*, and *liaX*); and transcription regulation (*tet*(M)).

**Figure 4.**
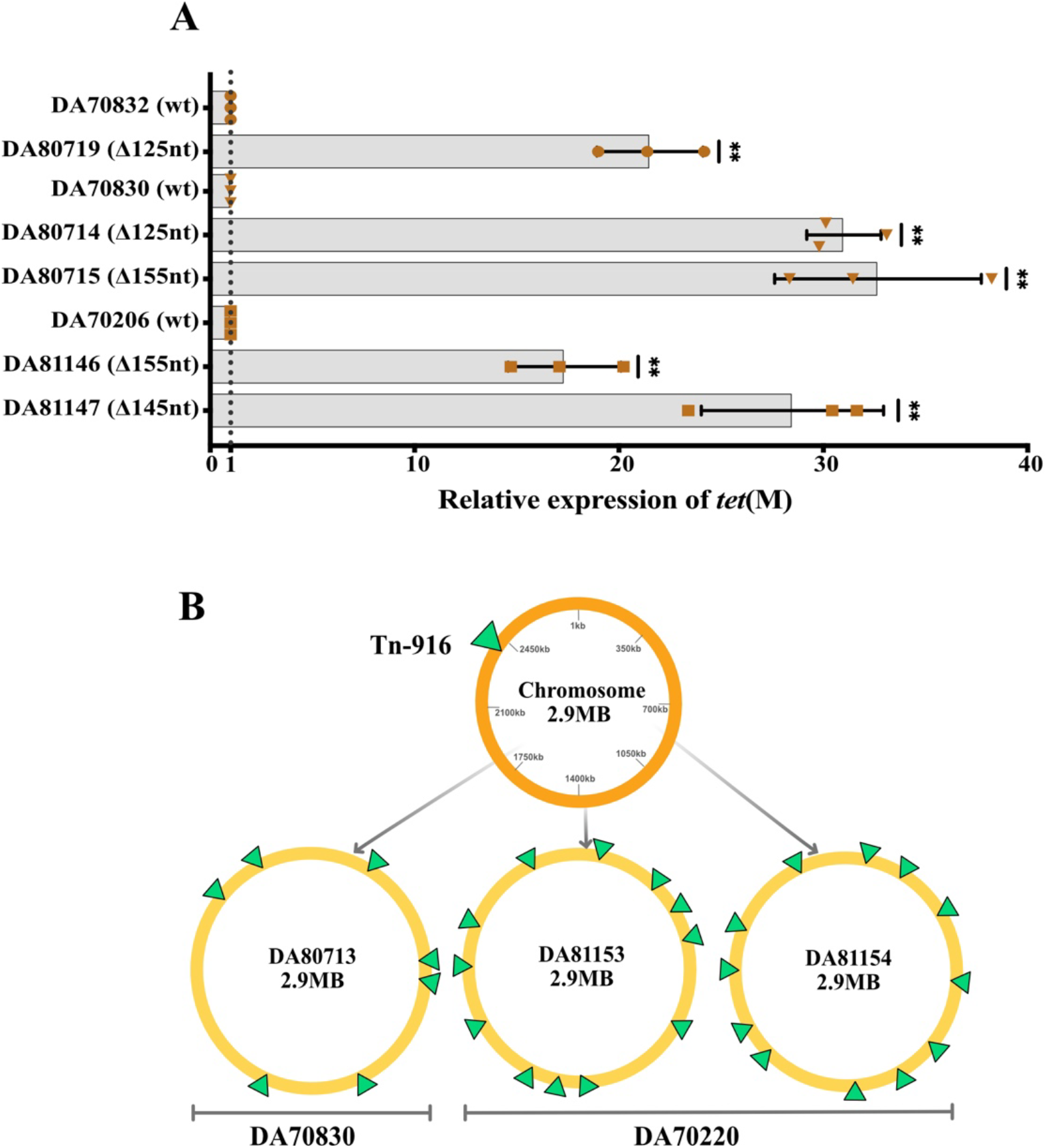
Fold increase in *tet*(M) expression and gene copies in TGC-resistant *E. faecalis* mutants. (A) RT–qPCR analysis of *tet*(M) expression in TGC-resistant mutants with a deletion in the leader peptide region 40 nucleotides upstream of *tet*(M). Expression is presented relative to that in the parental strains. The y-axis shows mutants names with deletion size in parentheses; parental isolates are labeled WT (wild type). Statistical significance was determined using a two-tailed t-test and is indicated as follows: *p* < 0.05 (*), *p* < 0.01 (**\*\***). Error bars represent standard deviation of n=3 biological replicates. The dashed line at 1 indicates baseline expression in parental strains. (B) Schematic representation of Tn916 transposition events in TGC-resistant mutants. Tn insertion sites (green triangles) on the chromosome of the mutants (yellow circles) or on the chromosome of the parental strains (red circle) are represented. DA numbers inside circles correspond to mutants with transposition events, and parental isolate for each mutant is indicated below each yellow circle. The Tn916 transposon carrying *tet*(M) was inserted into multiple chromosomal sites: six in DA80713, eleven in DA81153, and eleven in DA81154.

### Minimal cross-resistance and occasional collateral sensitivity in resistant mutants

Cross-resistance (CR) analysis was performed for resistant mutants that only carried a single point mutation in one gene to determine whether that mutation alone could confer resistance to other antibiotics. To assess whether the selected mutants exhibited CR to other drugs, we compared MIC values of resistant mutants with their corresponding parental isolates across seven additional antibiotics. Relative MIC fold-changes of these strains are summarized in Supplementary Fig. 5. Overall, resistant mutants exhibited no significant CR; most MIC changes were <2-fold, indicating minimal variation from parental strains. One GEN-resistant mutant with a *fusA* mutation showed a 2.6-fold increase in DAP MIC, while three DAP-resistant mutants with *liaF* and *liaS* mutations had 2-fold increases in the MIC of GEN. TGC-resistant mutants showed no consistent MIC increases for other antibiotics; 75% (10/28) of the TGC mutant-antibiotic combinations (corresponding to four isolates tested against seven antibiotics) had equal or lower MICs compared to parental isolates and no CR was detected for theses mutants (Fig. 4). In contrast, CS was observed in one DAP-resistant mutant (DA80691) toward AMX, GEN, and TGC, associated with a mutation in a DAK2 domain family protein, exceeding the >4-fold reduction in MIC relative to the parental isolate. CS was also detected in two GEN-resistant mutants (DA80707 and DA80709) with *rplF* deletions toward AMP, AMX and LNZ, respectively. No CS was observed among TGC-resistant mutants.

## Discussion

In this study, we found that HR was frequent for DAP, GEN, and TGC, but absent for LNZ and VAN (Fig. 1, and Table 2). Importantly, all HR isolates were classified as clinically susceptible by antimicrobial susceptibility testing (Etest).

Within the *Enterococcus* genus, where resistance to DAP remains below 2% (61), HR to DAP has rarely been reported. In our collection, the prevalence of DAP HR was 20%. One study identified HR in *E. faecium* isolated from patients, with 3 out of 6 isolates showing MIC variability in Sensititre assays, but PAP analysis was not performed to confirm the HR phenotype (31). For DAP, HR has previously been predominantly observed in methicillin-resistant *S. aureus* (MRSA), where prevalence rates of 7.4% (62) and 50% (63) have been reported. However, these studies employed either different methodologies or definitions: the first defined HR as growth of subpopulations at two-fold the clinical breakpoint, whereas the second applied the same PAP-based criteria as used in our study. These differences may account for the variation in the reported prevalence. A notably high prevalence of 25.6% of DAP HR, based on standardized PAP method, had been reported among clinical *S. aureus* isolates (8).

No comprehensive studies have examined GEN HR in *E. faecalis* prior to our study. In our collection, the prevalence of HR for GEN was 13.2%. The intrinsically low susceptibility of enterococci to aminoglycosides may explain the limited attention to HR in this genus. In contrast, HR to GEN is well-documented in Gram-negative bacteria (e.g., *Enterobacter cloacae* (64), *E. coli* (9), *Acinetobacter baumannii* (65), and *Klebsiella pneumoniae* (9,66)). Among Gram-positive bacteria, HR to GEN has been reported in *S. aureus*, with prevalence rates reaching up to 69.2% in clinical isolates using the same HR definition as we applied in this study, but using a different methodological approach (8).

Among the antibiotics tested, TGC had the highest prevalence of HR (35%) in our collection of isolates. This contrasts with a previous study where TGC HR had been detected in only 1.75% (7/399 isolates) *E. faecalis* clinical isolates (67). Studies on tetracycline derivatives such as omadacycline and eravacycline also reported relatively low HR prevalence frequencies of 2.5% and 8%, respectively for *E. faecalis* (34,36). The lower frequencies observed in these studies compared to ours may be attributed to differences in isolates origin or the criteria used to define HR. For example, the TGC-HR for 399 isolates in a previous study was defined as a TGC-susceptible isolate (MIC ≤0.25 mg/L) with subpopulations capable of growing at ≥0.5 mg/L TGC, using a detection threshold of 20 CFU/mL (67). These methodological differences likely affect reported prevalence and highlight the importance of standardized HR definitions in comparative studies.

No HR towards LNZ or VAN was detected among in our strain collection. Evidence for LNZ HR is scarce, largely case-based, and often restricted to *E. faecium* (76,77). High-level LNZ resistance typically requires G2576T substitutions in multiple copies of the 23S rRNA gene (31), which impair linezolid binding (68). The low probability of sequentially accumulating these multi-copy mutations likely explains the absence of LNZ-HR phenotypes in our isolate set.

To date, HR to VAN in *E. faecalis* has not been reported, with most existing data focusing on *E. faecium*. In enterococci, VAN HR has been attributed to either mutations in the *vanA* gene for the VanA resistance mechanism (39,69,70) or to tandem amplification of the genes encoding the VanM resistance mechanism (21,22). While we did not sequence all 40 isolates, in our 11 *E. faecalis* isolates exhibiting HR to DAP, GEN, or TGC, WGS showed that no *van* operons were present according to CARD and ResFinder databases (71). Thus, absence of VAN HR is probably due to a lack of *van* operon in our collection of isolates.

In our study, the primary mechanism underlying HR in *E. faecalis* involved mutations in chromosomal genes. While several of the mutated genes are known to confer resistance to specific antibiotics, others have not been previously described in Enterococcus species and may represent novel contributors to HR.

DAP resistance in enterococci primarily involved mutations in two key pathways: the *liaFSR* regulatory system, and genes linked to phospholipid metabolism (e.g., *gdpD*, *cls*). The *liaFSR* genes are well-characterized in *E. faecalis* and *E. faecium* where they mediate cell envelope stress responses and contributes to resistance against multiple antimicrobial peptides (55,72,73). In our study, ∼90% of the DAP-resistant mutants selected from HR isolates carried mutations in *liaFSR* or *liaX* encoding a sensor responsible for LiaFSR activation in presence of DAP (74). Unlike prior studies where Thr120Ala and Trp73Cys substitutions in *liaS* and *liaR*, respectively, were the most frequent alterations in *E. faecium* (75), we identified a Met142Leu substitution in *liaS* and a Leu171 duplication in *liaF*. In addition, frequent loss-of-function mutations in *liaX* which have previously been shown to contribute to DAP resistance in this genus (76,77) were also observed in our isolates. Thus, our findings highlight alterations in LiaFSR and LiaX as major causes of HR to DAP in *E. faecalis*. About 10% of DAP-resistant mutants selected had mutations in DUF4097 and DAK2 domain proteins, whose role in DAP resistance remain unclear.

Aminoglycoside resistance in enterococci typically involves reduced permeability, ribosomal target modification, or acquisition of aminoglycoside-modifying enzymes (29). However, in the context of HR, mechanistic insights in *E. faecalis* remain largely unexplored. In our study, nine mutants isolated from GEN HR isolates had mutations in the *rplF* gene. Mutations in *rplF*, which encodes for the ribosomal protein L6, are implicated in ciprofloxacin resistance in *Mycobacterium* (78) and GEN resistance in *E. coli* (57) and associated with a ≥2-fold increase in MICs of GEN in our study. Additionally, frequent mutations in *fusA* (elongation factor G) and *marR* genes were found. The role of *fusA*, encoding elongation factor G, in aminoglycoside resistance is well-documented in *E. coli* (57), *S. aureus* (8), and *Pseudomonas* spp (79), and for fusidic acid resistance in *E. faecium* (80). The functional overlap between fusidic acid and GEN, with both targeting the ribosomal elongation process, suggests that *fusA* mutations may also contribute to GEN resistance in *E. faecalis.* Mutations in *marR*, which encodes for a potential transcriptional regulator of the MarR family, were also selected in our seven GEN-HR mutants. In *E. coli*, *marR* regulates the multiple antibiotic resistance (*mar*) operon (81) and *marR* mutation decreases susceptibility to tetracyclines, quinolones and β-lactams, mainly through the up-regulation of the AcrAB-TolC multidrug efflux pump (82,83). However, *Enterococcus* lacks the canonical *marRAB* operon and the AcrAB-TolC pump, and how *marR* mutation increases MIC of GEN in *E. faecalis* remains unknown. However, recurrent isolation of *marR* mutations in our analysis suggests an important role of MarR in *E. faecalis*.

TGC resistance in *Enterococcus* has been primarily linked to alterations affecting the tetracycline-binding site on the 30S ribosomal subunit, including mutations in *rpsJ* (encoding ribosomal protein S10) and 16S rRNA (60), as well as overexpression of efflux pump regulators and ribosomal protection proteins such as *tet*(M)and *tet*(L) (37). In our study, 4 out of 17 TGC-resistant mutants selected from TGC HR isolates carried *rpsJ* mutations, with three carrying the same A54E substitution near the conserved KYKD motif (positions 57–60), a region implicated in TGC resistance (59,84). Indeed, diverse Gram-positive and Gram-negative species, including *E. faecalis*, have been shown to accumulate *rpsJ* mutations under TGC exposure (60), and Bai et al. also reported *rpsJ* and 16S rRNA variants in *E. faecalis* as associated with decreased TGC susceptibility, although they proposed efflux pump overexpression as the predominant HR mechanism (85). Although efflux-mediated TGC resistance is well documented in both Gram-positive and Gram-negative pathogens (86–88) and BMP-family ABC transporters were recently linked to tetracycline-derivative HR in *E. faecalis* against tetracycline derivatives (eravacycline, omadacycline) (34,36), no evidence of efflux involvement was detected in our TGC-HR isolates. The predominant mechanism of TGC HR in our study was a 125 to 155 bp deletion located 40 bp upstream of the *tet*(M) start codon, which was selected in 5/17 (29%) of the TGC-resistant mutants. This deletion disrupts the transcriptional attenuation stem-loop located 36 bp upstream of the start codon, leading to constitutive overexpression of *tet*(M) (89). Our findings are consistent with this regulatory model, as we observed a ≥25-fold increase in *tet*(M) expression in the five mutants carrying the upstream deletion compared to their parental isolates. A similar 125-bp deletion upstream of *tet*(M) had previously been described in *E. faecalis*, which led to a 10-fold increase in *tet*(M) expression (90), and comparable 87-bp deletions located 54 bp upstream of *tet*(M) have also been selected in TGC-adapted *S. aureus* (91). Three TGC-resistant mutants had Tn916-mediated transposition of *tet*(M) to multiple chromosomal locations (6–12 copies of Tn916 per TGC-resistant mutant), which increased *tet*(M) gene dosage and likely contributed to the elevated MIC of TGC (17). Tn916 is a broad-host-range conjugative transposon known to mediate *tet*(M) dissemination across enterococci and other Gram-positive species (92). Increased gene dosage of tetracycline-resistance genes through transposon of a *Tn*3 was also linked to TGC HR in *Klebsiella pneumoniae* (17). Finally, for five mutants, no genetic changes were detected despite repeated short read and Nanopore sequencing and reproducible increases in MICs of TGC, indicating possible epigenetic modifications.

The absence of tandem amplification-mediated increased gene dosage as a mechanism of HR in *E. faecalis* contrasts with Gram-negative bacteria, where they are major contributors to HR (9,16,19). One plausible explanation is the limited presence, in *E. faecalis*, of *bona fide* resistance genes that can confer resistance when present in higher copy numbers. WGS of our HR isolates confirmed that most lacked such genes which could give resistance to the antibiotic where they selected from, except DA70612 that carried *aad*(*6*), which is linked to streptomycin resistance in *E. faecalis* (93), rather than GEN resistance for which *aac(6′)-Ie-aph(2″)-Ia* is more relevant (29,94) (Supplementary Table 3). A second plausible explanation may be the absence of the genomic context required for tandem gene amplification, which typically involves homologous recombination between direct repeat sequences, such as IS elements, transposase genes, or rRNA operons, flanking the resistance gene. Indeed, sequence analysis of both the GEN-HR isolate DA70612 and the TGC HR isolates revealed a lack of such direct repeats. This reduces the probability of amplification events and might explain absence of *tet*(M) tandem amplifications despite presence of *tet*(M) in all four TGC HR isolates. In summary, the lack of tandem amplifications might reflect both (i) a low content of amplifiable resistance genes, particularly in isolates showing DAP and GEN HR, and (ii) absence of flanking repeats stimulating tandem amplification in TGC-HR isolates. In addition, *tet*(M) carriage was not specific to TGC-HR isolates, as this gene was also present in several other parental isolates (Supplementary Table 3). Why TGC HR by *tet*(M) mutation was observed in some *tet*(M)-harboring isolates but not others remain unknown. These findings, consistent with our prior *S. aureus* study (8), suggest that HR in Gram-positive bacteria may be mainly mutation-driven rather than mediated by increased resistance gene dosage.

Overall, most resistant mutants selected from HR isolates did not show CR, with only a few mutants displaying modest MIC increase (>2-fold) to other antibiotics. Mutations affecting *liaFSR* selected on DAP also showed increased GEN resistance, as previously described for *E. faecium* (72), and one GEN-resistant mutant with a *fusA* mutation had CR to DAP. Notably, CS was the predominant pattern in our dataset, particularly between aminoglycosides and β-lactams. This mirrors the findings of Wang et al. (95), who showed that AMX-resistant isolates were hypersensitive to GEN, while kanamycin-resistant isolates were more susceptible to β-lactams. They further demonstrated that *fusA* mutations, selected across multiple species including *E. faecalis*, are key drivers of CS. Indeed, two out of three *fusA* mutants that we selected displayed CS across several antibiotics, underscoring the evolutionary trade-offs of specific mutations (34).

This study provides a comprehensive analysis of HR in *E. faecalis*, showing its frequent occurrence for DAP, GEN, and TGC in clinical isolates classified as susceptible by conventional AST methods. HR in *E. faecalis* was caused by chromosomal mutational events including SNPs, deletions leading to overexpression of a resistance gene, or multiple transposition events increasing the gene copy number of a resistance gene. Importantly, HR was not caused by tandem amplification of resistance genes, distinguishing it from HR in Gram-negative bacteria. We found that resistance mutations often led to collateral sensitivity, particularly between aminoglycosides and β-lactams. We provide new insights in regard to HR in *E. faecalis* and advocate for the integration of HR detection into routine diagnostics.

## Supporting information

**S1 Fig. Frequency and MIC distribution among 40 clinical *E. faecalis* isolates for five antibiotics.** MICs were determined using Etests, and each dot represents the number of isolates (on the Y axis) sharing the same MIC value on the X-axis. The vertical dotted line indicates EUCAST clinical breakpoints for all antibiotics except GEN and DAP, where it represents the epidemiological cutoff value (ECOFF). The gray box shows the worldwide aggregated MIC distributions for each antibiotic according to EUCAST.

**S2 Fig. Distribution of HR and non-HR isolates of *E. faecalis* as a function of MIC value.** (ns) indicate nonsignificant differences between HR and non-HR groups in the Mann-Whitney test, with the *p*-values above 0.05. DAP (daptomycin), GEN (gentamicin), and TGC (tigecycline).

**S3 Fig. Fold-change in MIC values of resistant mutants derived from HR isolates**. Grey bars represent individual mutants, grouped by parental isolate. Black bars indicate the parental isolates, set to a relative MIC fold-change of 1 (shown by the horizontal dotted line), with corresponding mutants displayed immediately after. MIC values were determined using a single Etest (n=1).

**S4 Fig. Mutations identified in DAP (A), GEN (B), and TGC (C) resistant mutants.** Mutants are labeled with their DA number below the graph with their corresponding parental isolates indicated above the graph. The number of chromosomal insertions of transposon Tn916 containing the *tet*(M) gene is denoted as *n::(number)* for mutants harboring this mutation. Mutations are annotated as follows: *fs* = frameshift; *\** = stop codon; *Δ* = deletion; *Dup* = duplication.

**S5 Fig. Cross-resistance analysis of HR mutants against seven antibiotics.** The y-axis shows log_2_ fold changes in MIC values of individual mutants relative to their corresponding parental isolates. The x-axis shows the DA numbers of resistant mutants, each selected with the antibiotic indicated above the graph. Mutants showing a ≥2-fold increase in MIC for a given antibiotic are highlighted in red, indicating potential cross-resistance, whereas those showing a >4-fold decrease is highlighted in green, indicating collateral sensitivity.

**S1 Table. Antibiotic concentrations used for the selection of *E. faecalis* resistant mutants in the population analysis profile (PAP) assay.** For each parental isolate, 3–6 resistant clones were independently isolated following exposure to the corresponding antibiotic at **8× the MIC (mg/L)** of the main population.

**S2 Table. Oligonucleotide primers used for qRT-PCR.**

**S3 Table. Antibiotic resistance genes identified in heteroresistant *E. faecalis* isolates used for mutant selection and whole-genome analysis.** Resistance genes potentially conferring reduced susceptibility to the indicated antibiotics were identified using the Comprehensive Antibiotic Resistance Database (CARD) and ResFinder. Dark grey cells indicate the antibiotic used for mutant selection from the corresponding parental isolate.

## Acknowledgments

We would like to thank Örjan Samuelson (University Hospital North Norway, Tromsö, Norway), Niels Frimodt-Møller (Rigshospitalet, Copenhagen, Denmark), Volkan Özenci (Karolinska University Hospital, Solna, Sweden), and Fernando Baquero (Hospital Ramon y Cajal, Madrid, Spain) for providing us with the *E. faecalis* clinical isolates.

## Author Contributions

**Conceptualization:** Dan I. Andersson, Sarah Satola

**Formal analysis:** Sheida Heidarian, Karin Hjort

**Investigation:** Sheida Heidarian, Karin Hjort, Sarah Lohsen; Sarah Satola; Andrei Guliaev

**Methodology:** Sheida Heidarian; Sarah Lohsen; Sarah Satola; Andrei Guliaev; Karin Hjort

**Resources:** Dan I. Andersson; Sarah Satola

**Supervision:** Karin Hjort, Hervé Nicoloff, Dan I. Andersson

**Writing – original draft:** Sheida Heidarian

**Writing – review & editing:** Sheida Heidarian, Andrei Guliaev, Sarah Satola, Sarah Lohsen, Karin Hjort, Hervé Nicoloff, Dan I. Andersson

