## Supplementary material for "Heteroresistance in *Enterococcus faecalis* is prevalent for key antibiotics and mainly caused by mutations": Supplemmentary figure

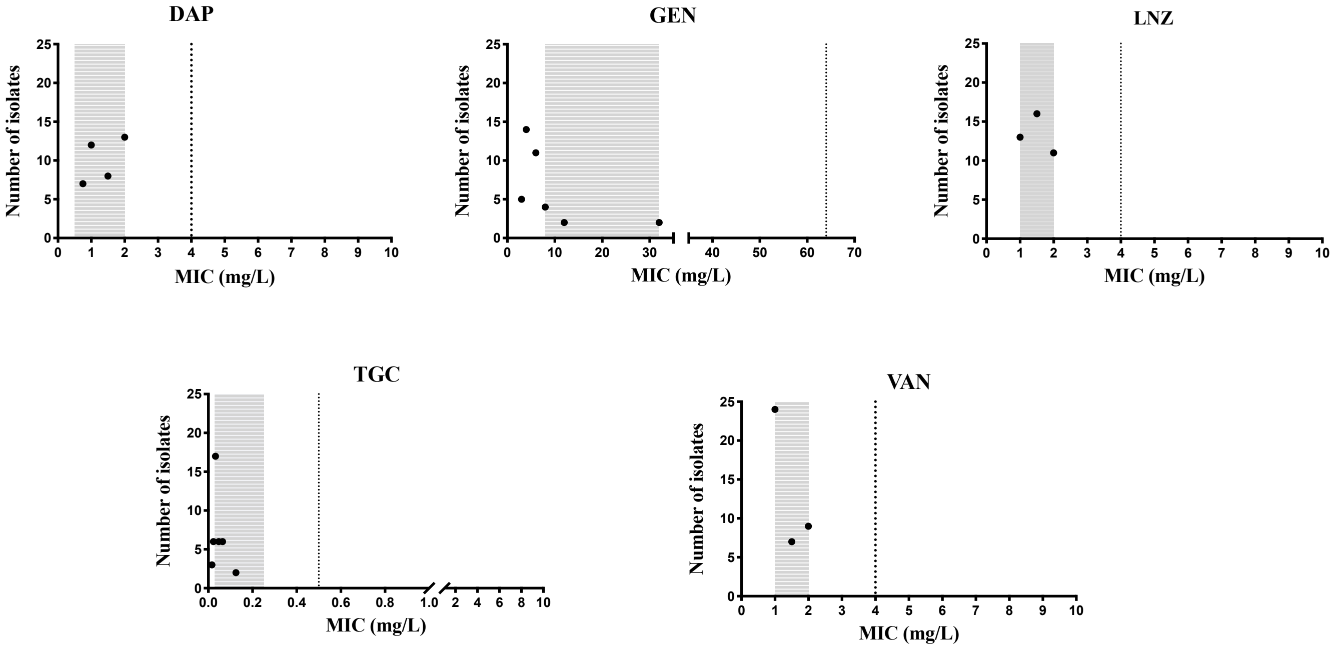


**Supplementary Fig 1. Frequency and MIC distribution among 40 clinical *E. faecalis* isolates for five antibiotics.** MICs were determined using E-tests, and each dot represents the number of isolates (on the Y axis) sharing the same MIC value on the X-axis. The vertical dotted line indicates EUCAST clinical breakpoints for all antibiotics except GEN and DAP, where it represents the epidemiological cutoff value (ECOFF). The gray box shows the worldwide aggregated MIC distributions for each antibiotic according to EUCAST.

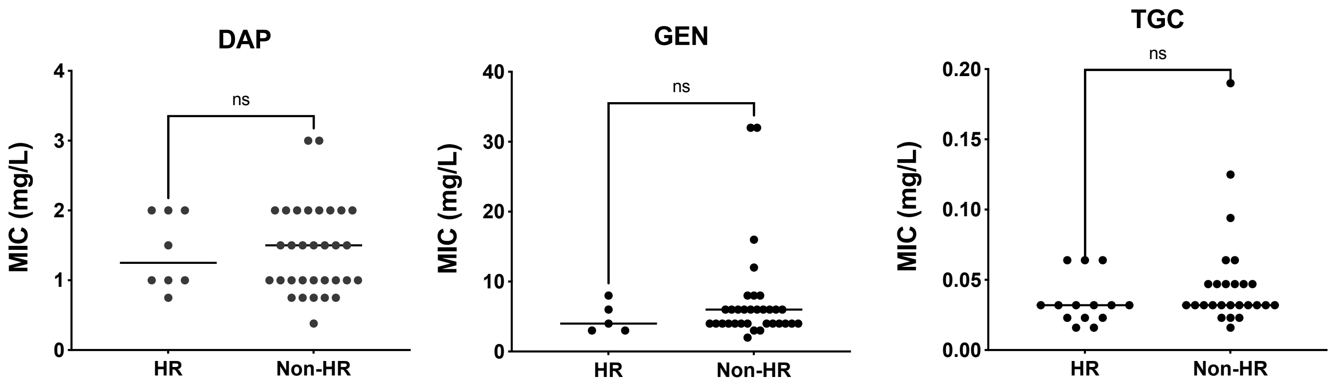


**Supplementary Fig 2. Distribution of HR and non-HR isolates of *E. faecalis* as a function of MIC value.** (ns) indicate nonsignificant differences between HR and non-HR groups in the Mann-Whitney test, with the *p*-values above 0.05. DAP (daptomycin), GEN (gentamicin), and TGC (tigecycline).

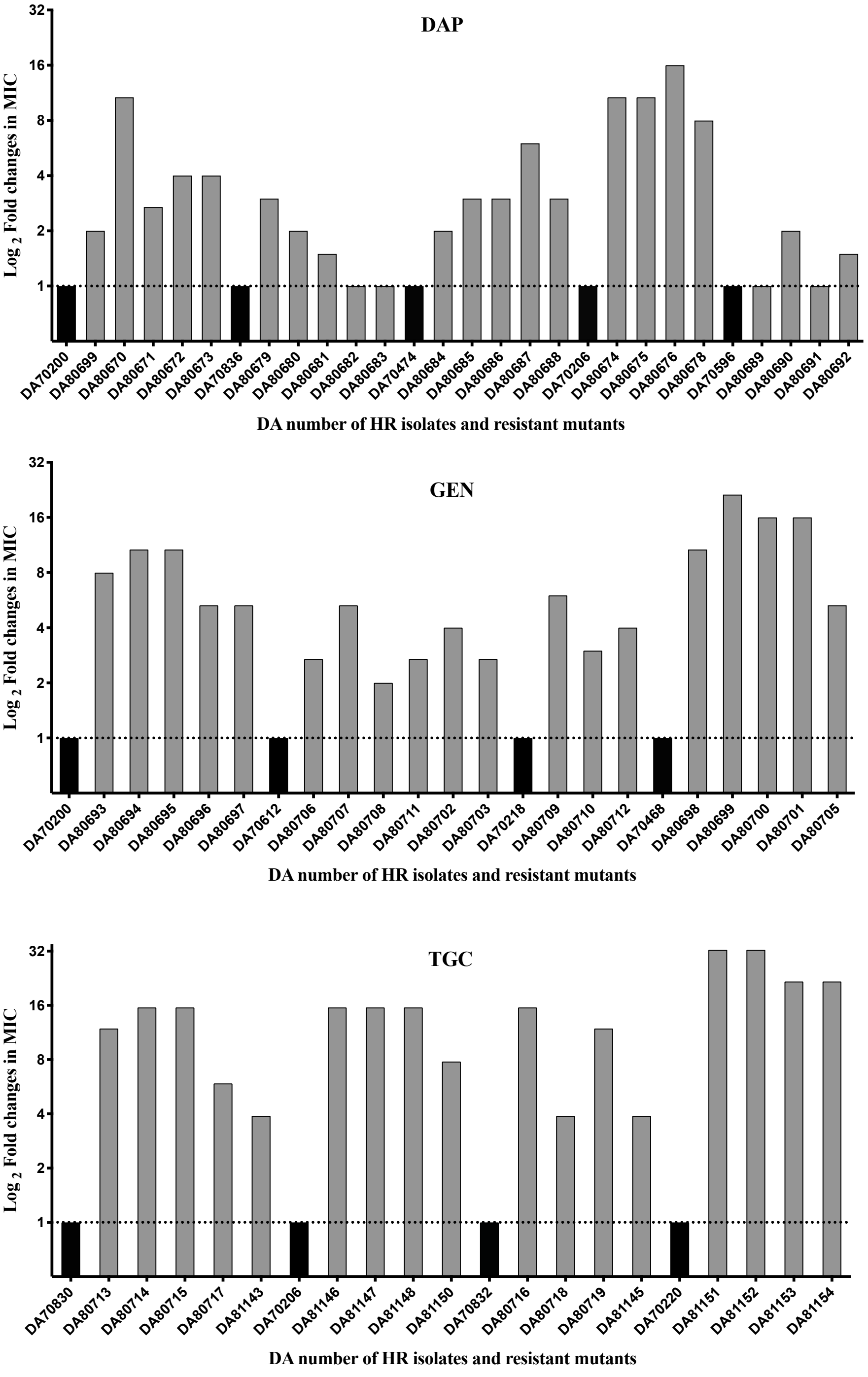


**Supplementary Fig 3. Fold-change in MIC values of resistant mutants derived from HR isolates**. Grey bars represent individual mutants, grouped by parental isolate. Black bars indicate the parental isolates, set to a relative MIC fold-change of 1 (shown by the horizontal dotted line), with corresponding mutants displayed immediately after. MIC values were determined using a single Etest (n=1).

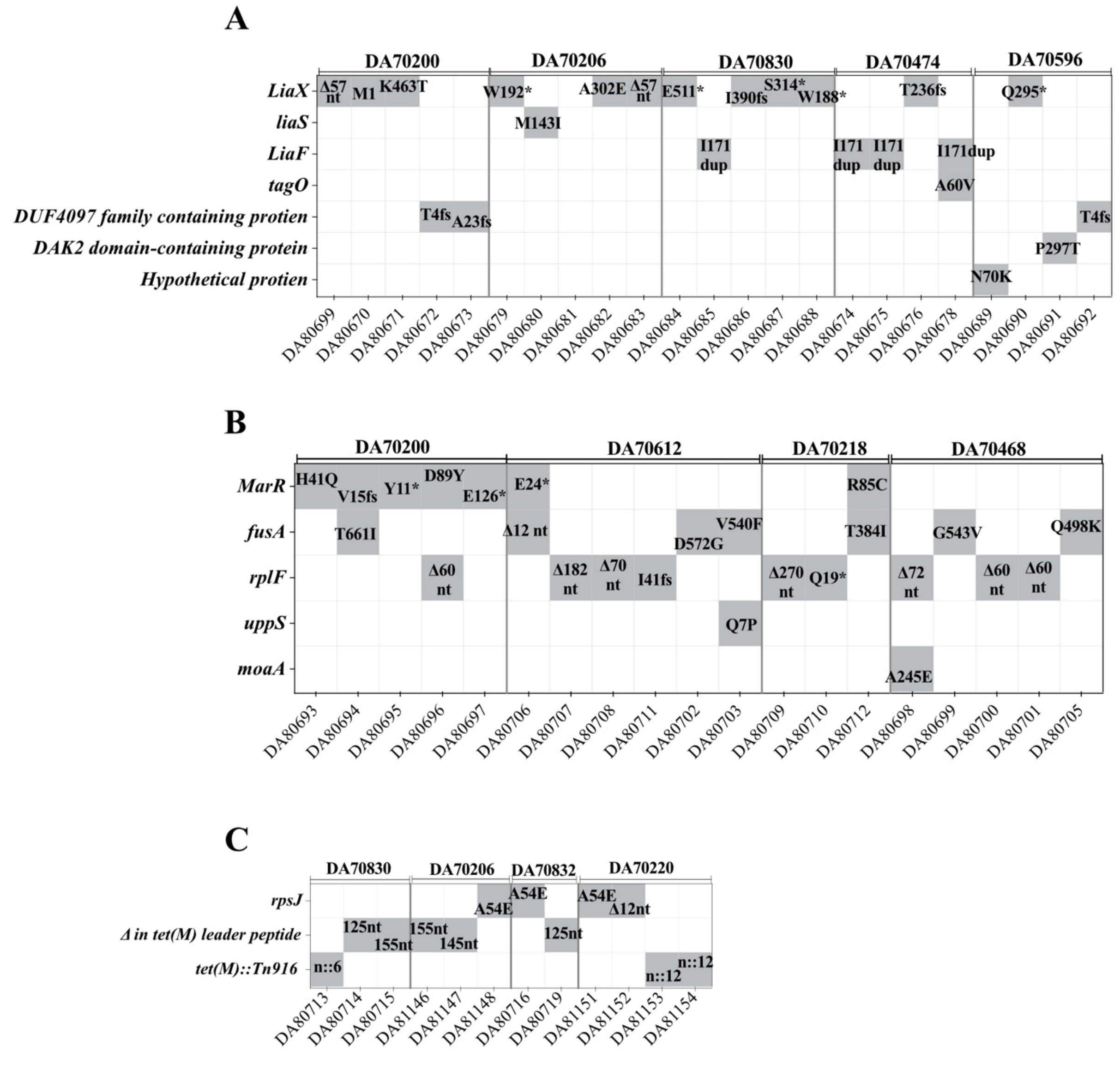


**Supplementary Fig 4. Mutations identified in DAP (A), GEN (B), and TGC (C) resistant mutants.** Mutants are labeled with their DA number below the graph with their corresponding parental isolates indicated above the graph. The number of chromosomal insertions of transposon Tn916 containing the *tet*(M) gene is denoted as *n::(number)* for mutants harboring this mutation. Mutation are annotated as follows: *fs* = frameshift; * = stop codon; *Δ* = deletion; *Dup* = duplication.

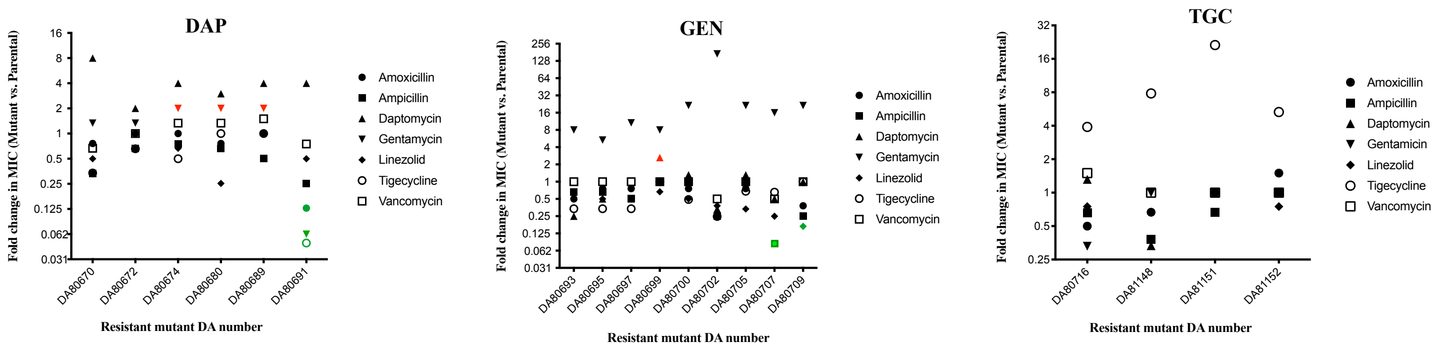


**Supplementary Fig 5. Cross-resistance analysis of HR mutants against seven antibiotics.** The y-axis shows $log_2$ fold changes in MIC values of individual mutants relative to their corresponding parental isolates. The x-axis shows the DA numbers of resistant mutants, each selected with the antibiotic indicated above the graph Mutants showing a ≥2-fold increase in MIC for a given antibiotic are highlighted in red, indicating potential cross-resistance, whereas those showing a >4-fold decrease is highlighted in green, indicating collateral sensitivity.

**Supplementary Table 1. Antibiotic concentrations used for the selection of E. faecalis resistant mutants in the population analysis profile (PAP) assay.** For each parental isolate, 3–6 resistant clones were independently isolated following exposure to the corresponding antibiotic at 8× the MIC (mg/L) of the main population.

| Resistant mutant ID | Parental isolate | Antibiotic used for selection | MIC (mg/L) of main population | Antibiotic concentration used for selection (8×MIC, mg/L) |
|---|---|---|---|---|
| DA80699<br>DA80670<br>DA80671<br>DA80672<br>DA80673 | DA70200 | DAP | 1.5 | 12 |
| DA80674<br>DA80675<br>DA80676<br>DA80678 | DA70206 | DAP | 0.75 | 6 |
| DA80679<br>DA80680<br>DA80681<br>DA80682<br>DA80683 | DA70836 | DAP | 2 | 16 |
| DA80684<br>DA80685<br>DA80686<br>DA80687<br>DA80688 | DA70474 | DAP | 2 | 16 |
| DA80689<br>DA80690<br>DA80691<br>DA80692 | DA70596 | DAP | 2 | 16 |
| DA80693<br>DA80694<br>DA80695<br>DA80696<br>DA80697 | DA70200 | GEN | 3 | 24 |
| DA80698<br>DA80699<br>DA80700<br>DA80701<br>DA80705 | DA70468 | GEN | 3 | 24 |

| | | | | |
|---|---|---|---|---|
| DA80702<br>DA80703<br>DA80706<br>DA80707<br>DA80708<br>DA80711 | DA70612 | GEN | 6 | 48 |
| DA80709<br>DA80710<br>DA80712 | DA70218 | GEN | 4 | 32 |
| DA80713<br>DA80714<br>DA80715<br>DA80717<br>DA81143 | DA70830 | TGC | 0.032 | 0.256 |
| DA80716<br>DA80718<br>DA80719<br>DA81145 | DA70832 | TGC | 0.032 | 0.256 |
| DA81146<br>DA81147<br>DA81148<br>DA81150 | DA70206 | TGC | 0.032 | 0.256 |
| DA81151<br>DA81152<br>DA81153<br>DA81154 | DA70220 | TGC | 0.023 | 0.184 |

**Supplementary Table 2. Oligonucleotide primers used for qRT-PCR**.

| Primer name | Sequence (5′→3′) | Target Gene | Amplicon Size (bp) |
|---|---|---|---|
| tet(M)_F<br>tet(M)_R | CTGTATCACCGCTTCCGTTG<br>CAGTCCGTCACATTCCAACC | *tet*(M) | 148 |
| gdh_F<br>gdh_R | AGATGCTGAAGTGATGCGCT<br>ACCAATTTCACGAGCGCCTA | *gdh* | 115 |

**Supplementary Table 3. Antibiotic resistance genes identified in heteroresistant *E. faecalis* isolates used for mutant selection and whole-genome analysis**. Resistance genes potentially conferring reduced susceptibility to the indicated antibiotics were identified using the Comprehensive Antibiotic Resistance Database (CARD) and ResFinder. Dark grey cells indicate the antibiotic used for mutant selection from the corresponding parental isolate.

| Parental isolate | DAP | GEN | TGC |
|---|---|---|---|
| DA70200 | | | *tet*(M) |
| DA 70206 | | | *tet*(M) |
| DA 70218 | | | *tet*(M), *tet*(L) |
| DA70220 | | *AAC(6')-Ie-APH(2")-Ia, aad(6)* | *tet*(M) |
| DA 70468 | | | |
| DA 70474 | | | |
| DA 70596 | | | *tet*(M) |
| DA 70612 | | *aad(6)* | *tet*(M) |
| DA 70830 | | | *tet*(M) |
| DA 70832 | | | *tet*(M) |
| DA 70836 | | | *tet*(M) |
